# Children’s Species Literacy as Estimated and Desired by Biodiversity Communicators: a Mismatch with the Actual Level

**DOI:** 10.1101/2021.11.10.466733

**Authors:** Michiel J. D. Hooykaas, Cathelijn Aten, Elisabeth M. Hemelaar, Casper J. Albers, Menno Schilthuizen, Ionica Smeets

## Abstract

While biodiversity decline continues and laypeople’s knowledge about species is limited, especially in children, high-quality communication is needed to raise awareness. For this, communicators should be aware of current knowledge levels in their target groups. We compared biodiversity communicators' estimates of the average species literacy level in primary school children with the actual level. Moreover, we explored the importance that communicators placed on species literacy and the level that they desired. Estimations of children’s average knowledge level varied widely and differed from the actual level. In particular, communicators overestimated the species literacy level. Although most biodiversity communicators agreed that knowledge about species is important, their view differed as to why species literacy would be important. Moreover, communicators differed with respect to the relative importance attached to different knowledge components. Professionals may thus benefit from a detailed framework of species literacy that illustrates different aspects and values. Most importantly, our findings suggest that to bridge the gap between actual and desired knowledge levels in children effectively, biodiversity communicators first need to become more aware of current perceptions in young audiences.

## 1. Introduction

At a time of great biodiversity loss and a widening gap between people and nature, conservationists are faced with a challenging task to build broad-based support for conservation (Ceballos et al., 2015, 2017; Miller, 2005; Pyle, 2011). Communicators can make a valuable contribution by raising awareness about biodiversity in the public (Bickford et al., 2012). However, while certain segments of society have successfully been reached, it has been acknowledged that, overall, laypeople are not well-informed about biodiversity (Navarro-Perez and Tidball, 2012), showing that communication about biodiversity has not yet been as effective as it could be.

Studies in different countries have demonstrated that laypeople, particularly primary school children, lack broad as well as in-depth knowledge about species (Balmford et al., 2002; Huxham et al., 2006; Torkar, 2016); i.e., they have low levels of species literacy (Hooykaas et al., 2019). For instance, in the Netherlands primary school children regularly failed at identifying common, native animals that can be easily encountered (Hooykaas et al., 2019), implying that they are disconnected from their local environment. This indicates that barriers need to be overcome by biodiversity communicators, as unknown species will not easily strike a chord with the public and their names may be perceived as jargon.

For biodiversity communicators it is important to take into account the knowledge levels present in their audiences, as these influence people’s expectations and determine the ways they will respond (Buijs et al., 2008; Thompson and Zamboanga, 2003). Prior knowledge affects subsequent learning and plays an important role in the construction of new understanding (Hailikari et al., 2007, 2008; National Research Council, 2000, 2007, 2009). To achieve high-quality communication, communicators should therefore connect to people’s knowledge base in a strategic manner. Messages will then be better comprehended and more readily received, and learning outcomes will be more likely to be in line with those intended (Wratten and Hodge, 1999).

However, before communicators can craft messages or devise strategies according to people’s existing knowledge, they should first be aware of it. It is therefore imperative that they can accurately estimate knowledge levels in their audiences. Yet, studies conducted outside of the field of biodiversity communication have demonstrated that estimating prior knowledge can be quite hard. For example, nursing professionals and physicians regularly experience difficulties in estimating health literacy in their patients (Bass et al., 2002; Kelly and Haidet, 2007; MacAbasco-O’Connell and Fry-Bowers, 2011), frequently resulting in overestimations (Dickens et al., 2013). In addition, teachers have been reported to fail at accurately estimating knowledge levels in their students (Perrenet, 2010; Schutte, 2010; Storm, 2012).

A mismatch between estimated and actual knowledge levels poses a problem as it may hamper communication. Overestimations can lead communicators to calibrate their language to a level above that of their public, resulting in messages that are not understood correctly by the audience, while underestimations may lead to needless repetition of information (Kelly and Haidet, 2007; Schutte, 2010). For instance, nature guides or text editors unaware of low species literacy levels may mention species names that act as jargon, while those who underestimate knowledge levels may elaborate on already well-known species, which may bore people and will not expand their perceptions of biodiversity. Ultimately, a bad fit may prevent educational and communicational goals from being achieved (Bass et al., 2002; Hailikari et al., 2008); e.g., it could make it harder to foster species literacy effectively and could hamper citizen science projects where participants are asked to count and record species (Falk et al., 2019).

Although research on knowledge estimations has been conducted in other fields of expertise, such as healthcare and education, no previous study has investigated biodiversity communicators’ perceptions of knowledge levels in laypeople. Research in this direction is important, as it may help explain current communication outcomes and can aid biodiversity communicators in reaching out successfully to broader audiences than before, so that eventually broad-based support for biodiversity conservation can be realized. It is especially relevant to study communicators’ awareness of knowledge levels in primary school children, as they are at a suitable age to learn about species and represent a generation that holds the key in addressing the biodiversity crisis in the future (Kahn Jr., 2002; Kellert, 1985, 2002; Magntorn and Helldén, 2006; White et al., 2018).

In addition to accurate estimations of knowledge levels in their audiences, communicators benefit from having a clear picture of what level of knowledge they strive for in their audiences. This can help set educational goals and provide clarity about the steps needed to achieve desired outcomes. While biodiversity communicators are expected to regard knowledge about biodiversity valuable and important, it is not yet clear what their views are about specific forms of it, such as species literacy. For instance, it is not known what the desired levels of species literacy would be and if and why communicators think that knowledge about species is important or not. Research in this direction can provide insight into the values attached to knowledge about biodiversity, and biodiversity communicators, educators, and conservationists may use this information to underline the importance of their own activities.

In this study we compared the average species literacy level of primary school children as estimated by biodiversity communicators in the Netherlands with the actual level, which had been determined during a previous project carried out just before the current study (Hooykaas et al., 2019). We further compared the estimated and actual average species literacy levels with the desired level, and we explored the importance placed by biodiversity communicators on species literacy.

We investigated the following research questions:

1. Are biodiversity communicators aware of the species literacy level in primary school children aged 9-10 years old?
2. What is the desired level of species literacy in primary school children aged 9-10 years old according to biodiversity communicators and how does this compare to the actual level?
3. What importance do biodiversity communicators place on species literacy in laypeople?

## 2. Methods

We constructed a survey (Appendix A) in Qualtrics (https://www.qualtrics.com) targeted at Dutch biodiversity communicators: people who communicate nature, biodiversity or animals in their voluntary or paid work. The survey was administered between May and July 2018, by sending an invitation via e-mail to a large number of Dutch organizations and institutions involved with nature and biodiversity, such as nature conservancy organizations, environmental education institutions, ecological consultants, and zoos. Participation was anonymous, avoiding social desirability or ‘prestige bias’ in the answers and taking into account privacy regulations (Streiner, David et al., 2015).

First, the communicators were asked to take a species identification test that had just been used during a different part of an overarching research project on communicating biodiversity, to assess species literacy levels in Dutch primary school children aged 9-10 years old. Full methods are described in Hooykaas et al. (2019). The identification test comprised 27 animal species native to the Netherlands, and participants were asked to provide the name of each depicted species, thereby identifying it as precisely as possible. Included species were mainly those occurring regularly in Dutch (sub)urban areas (e.g., house sparrow (*Passer domesticus*)), supplemented by a few species encountered predominantly outside urban areas (e.g., wild boar (*Sus scrofa*)). In the test, each animal was represented by one or two color pictures from the website https://pixabay.com/ – see Figure 1.

**Fig. 1.**
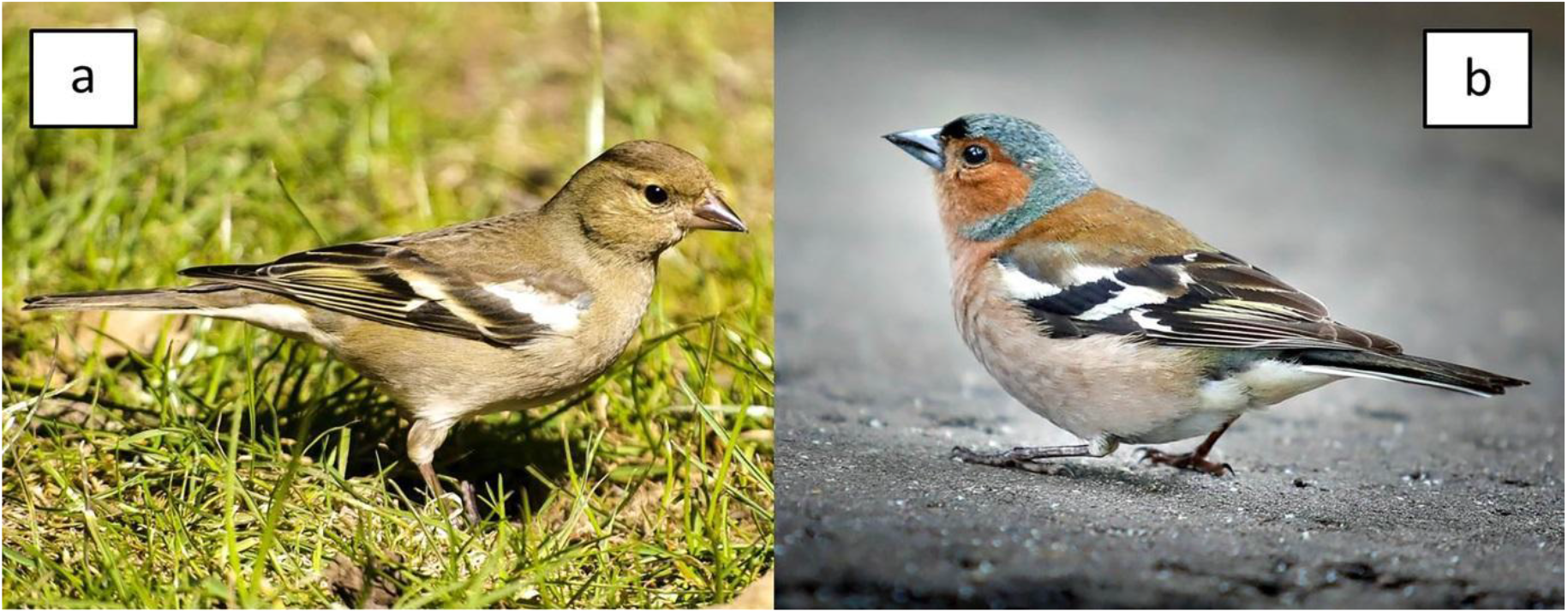
Female (a) and male (b) chaffinch (*Fringilla coelebs*); photo credits a. [Kathy Büscher](https://pixabay.com/nl/photos/vink-vrouwen-vink-vink-vogel-vogel-2404776/) b. [Klimkin Sergey](https://pixabay.com/nl/photos/vink-vogel-mus-het-park-een-2491484/).

After communicators had finished the species identification test, they were asked to estimate the species literacy level of primary school children aged 9 or 10 years old (i.e. their average achieved identification score: the number of correct identifications), and they were asked what the desired species literacy level in this group would be (i.e. the desired average achieved identification score). Communicators were also asked whether or not they had targeted primary school children aged 9-10 in their communication in the past 5 years, to investigate the influence of experience with the target group on estimation accuracy. Finally, we explored the importance placed by biodiversity communicators on species literacy, by asking them whether they agreed with the statement “it is important for people to recognize many animal species” on a 10-point scale and offering them the possibility to elaborate their answer with arguments.

### 2.1. Analyses and statistical procedures

Data were compiled in Microsoft Excel and subsequently processed with IBM SPSS Statistics 25.0. First, we used Welch’ independent samples *t*-tests to compare the average species literacy level in primary school children aged 9-10 as estimated and considered desirable by the communicators on the one hand with the actual level on the other. For the actual species literacy level, we used the average achieved identification score of 602 children (M = 9.5, SD = 3.4), established during the research project mentioned before that took place just prior to the current project; most children (86.9%) had recognized less than half of the species. Moreover, we compared the communicator-estimated average species literacy level in primary school children aged 9-10 by the communicators with the level considered desirable using a paired *t*-test. To account for multiple testing, a strict Bonferroni correction was applied.

To provide insight into the importance placed by biodiversity communicators on species literacy, we analyzed the answers to the 10-point scale question, and we used pattern analysis (Braun and Clarke, 2006) to carry out inductive coding of the additional remarks provided by the participants. The codes were eventually grouped into categories. To avoid subjectivity, codes and categories were designed by three researchers and discussed among colleagues. Depending on the variation in arguments provided by the participants, each answer received one or more codes (identical codes were not repeated). After one researcher had coded the dataset, half of the coded answer fragments were selected randomly and coded independently and blind to the previous coding by a second researcher. Intercoder reliability was high (percent agreement = 81%, Cohen’s Kappa = 0.798), indicating a strong level of agreement between the two coders (McHugh, 2012). Subsequently, the discrepancies were discussed by the coders and resolved.

## 3. Results

### 3.1. Descriptive statistics

The final dataset (Appendix B) included 677 biodiversity communicators (e.g., nature guides, communicators in zoos, spokespersons and text editors at nature conservancy organizations, and ecological consultants).

### 3.2. Species literacy estimations by communicators

Communicators’ estimations of the average species literacy level in primary school children aged 9-10 varied widely and regularly differed from the actual level – see Figure 2. The average identification score in primary school children as estimated by communicators (M = 11.4, SD = 4.2) was higher than the actual achieved score in this group (M = 9.5, SD = 3.4); *t*(1269.5) = 9.20, *p* < .001. In fact, 53.5% of the communicators overestimated the knowledge level (e.g., one in three incorrectly assumed that the average child would correctly identify over half of the species). Only one in four communicators (25.0%) estimated species literacy in children accurately, at an average achieved identification score of 9 or 10 out of 27 species, and 21.6% of the communicators underestimated species literacy in primary school children.

**Fig. 2.**
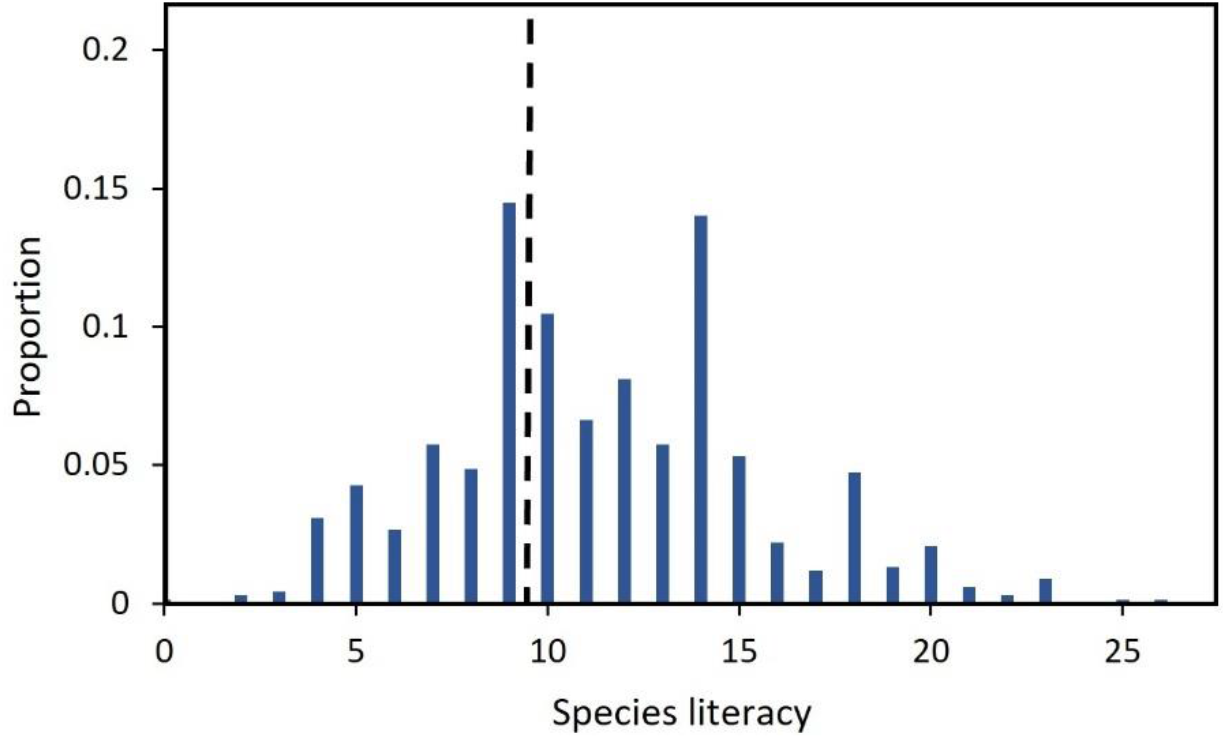
Distribution of biodiversity communicators’ estimations of the average species literacy level (i.e. identification score) in primary school children aged 9-10. The actual level, established during a previous research project just prior to the current study, is depicted with a dashed line. We note that communicators were asked to estimate the species literacy level on a scale from 0 to 27, where a few levels (e.g., 5, 9, 14) were indicated. Although this may explain the peak at 9 species, and might thus have increased the number of communicators with accurate estimations, the wide range in estimations demonstrates clearly that most communicators were unaware of the actual knowledge level.

Next, we investigated the influence of experience with primary school children as a target group on communicators’ estimations, by comparing the estimates of children’s species literacy made by communicators with (59.8%) and without (40.2%) children aged 9-10 as a target group. Estimations by communicators with children as a target group (M = 11.4, SD = 4.2) and by communicators without children as a target group (M = 11.5, SD = 4.1) did not differ significantly, *t*(589.67) = 0.34, *p* = .736).

### 3.3. Desired levels of species literacy

To further put children’s species literacy level in perspective, we compared the actual and estimated level with the level as desired by the communicators. Significant differences were found. The desired average species literacy level (M = 14.8, SD = 5.1) was considerably higher than both the actual average level (M = 9.5, SD = 3.4); *t*(1197.1) = 22.11, *p* < .001 and the estimated average level (M = 11.4, SD = 4.2); *t*(676) = 19.39, *p* < .001. While 23.3% of the communicators would be satisfied with the actual species literacy level (desiring no more than 10 out of 27 species to be correctly identified), the majority (76.7%) wished for a higher knowledge level – see Figure 3. For instance, two in three communicators (65.9%) expressed that children should be able to identify over half of the species.

**Fig. 3.**
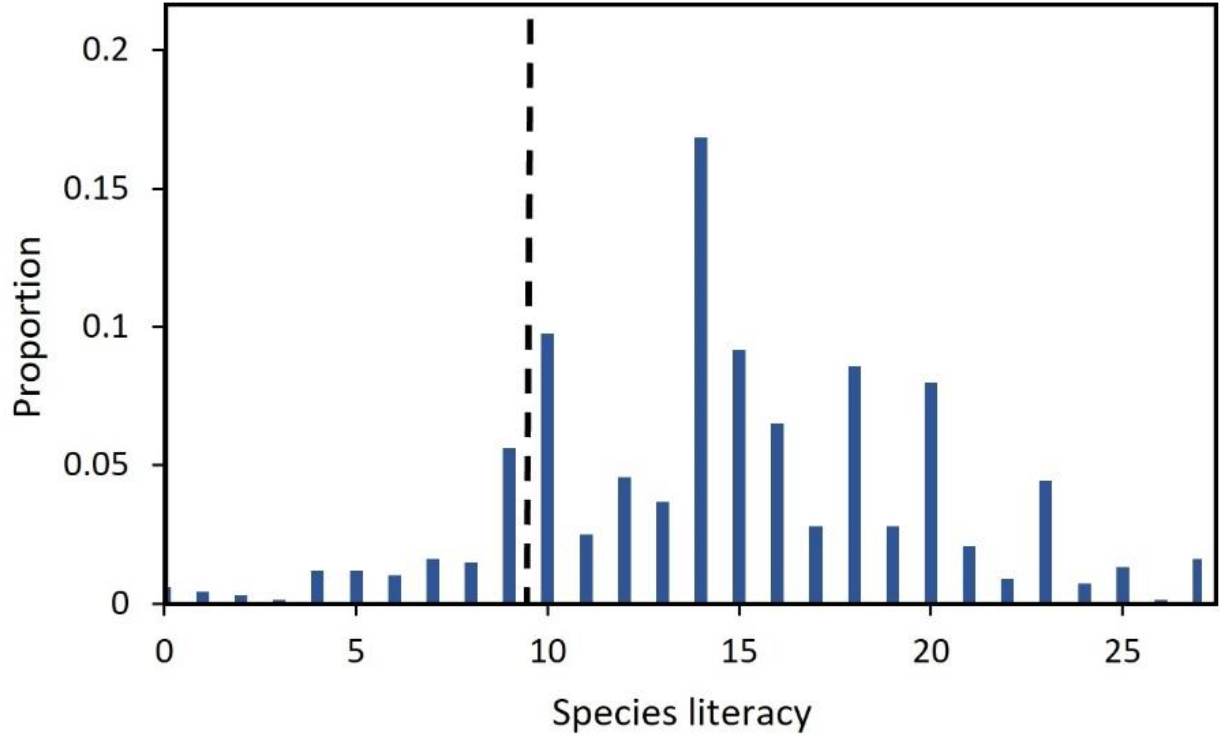
Distribution of the desired average species literacy level (i.e. identification score) in primary school children aged 9-10 according to biodiversity communicators. The actual level, established during a previous research project just prior to the current study, is depicted with a dashed line.

### 3.4. Importance placed on species literacy

The majority of the communicators attached importance to species literacy; on a 10-point scale 78.7% provided scores of 6 to 10 to the statement that people should be able to recognize many different animal species. Only a minority of the participants (4.9%) placed little to no importance on knowledge about species in laypeople (score 0 to 4).

To provide further insight into communicators’ perceptions of the importance of species literacy, we carried out inductive coding of the remarks provided by the participants. Each answer received 1 or more codes, and the total number of coded answer fragments (634) exceeded the number of communicators that provided remarks (439 out of 677). There were seventeen different codes grouped into three categories: *1 = Species literacy is important,* 2 = *Species literacy is not important*, and 3 = *Species literacy is not as or as important as…* – see Table 1. Each category contained the same four themes (insight, interest/experience, affinities/care, well-being) supplemented by a few separate codes. In addition, an eighteenth code contained 69 fragments that could not be assigned any of the previous 17 codes, e.g., because they were not an answer to the actual question (‘*the more knowledge, the better*’) or neutral (‘*no opinion*’).

**Table. 1.**
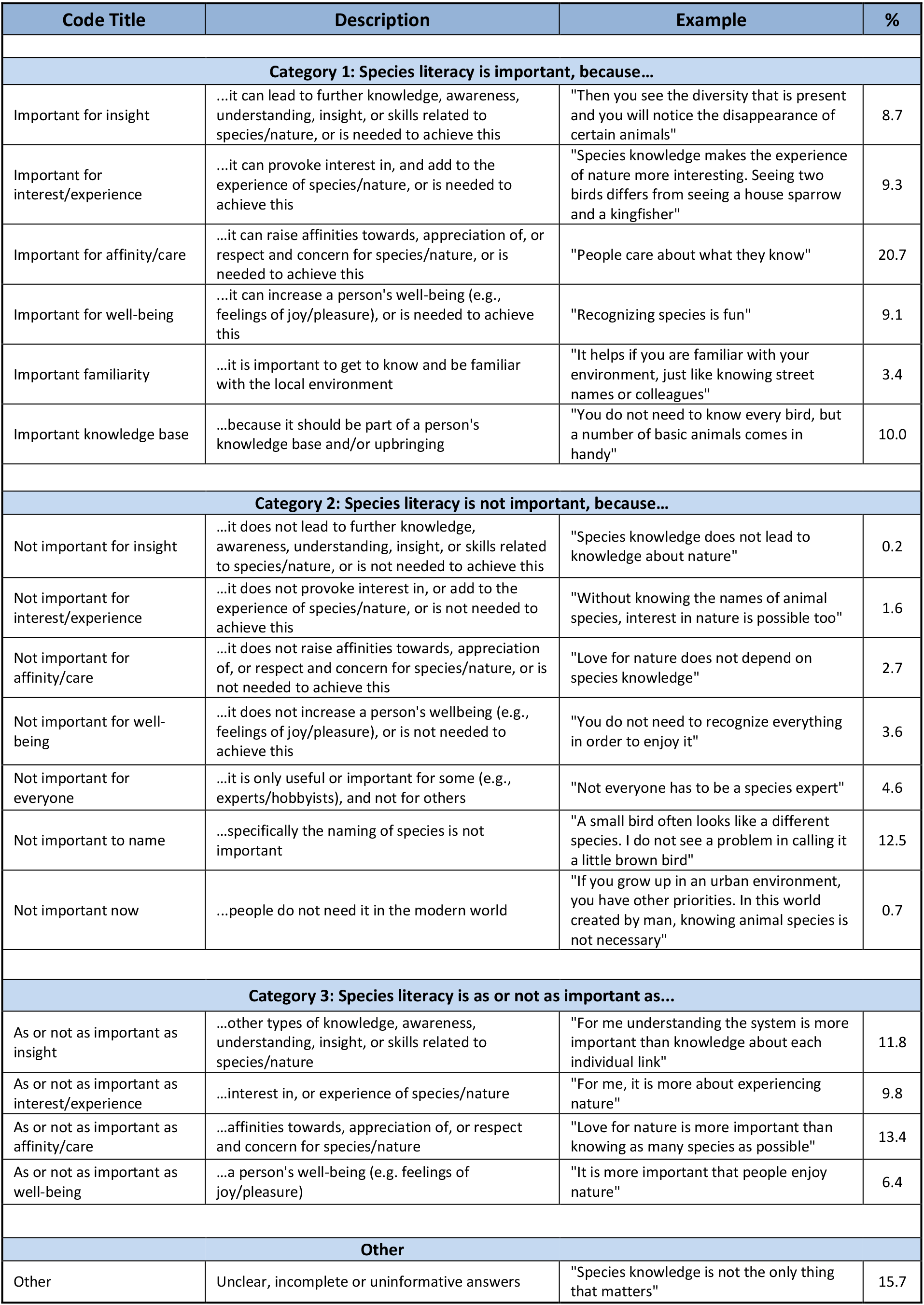
Overview of the codes and categories used during the inductive coding process of the remarks made by the communicators. The percentages show how many of the 439 communicators providing remarks used an argument with that particular code.

Different reasons were expressed by the biodiversity communicators as to why knowledge about species would be important or not. Of the coded answer fragments, 42.4% underlined the importance of species literacy. In particular, a considerable number of communicators expressed that species knowledge may help to create affinities towards nature and species, ultimately contributing to conservation. Participants also argued that knowledge about species, common everyday species especially, should be part of any person’s knowledge base, in line with comments from communicators that it is important specifically to be familiar with your surroundings. Furthermore, communicators noted that knowledge about species can provoke curiosity and can strengthen nature experiences, can contribute to well-being, e.g., by triggering joy and building a person’s confidence to talk about nature, and that knowledge and skills related to species (e.g., observing) can lead to further insights and broader understanding. For example, people knowledgeable about species may notice and pay attention to ongoing changes in population densities.

Of the coded answer fragments, 18% were objections against the idea that species literacy would be important. For instance, some communicators considered knowledge about species to be useful only for experts and hobbyists and a few expressed that people nowadays do not need knowledge about species, because information can be retrieved quickly and citizens are less directly dependent on natural resources. In particular, we found evidence for a lack of agreement among professionals of the importance of knowing species names; it was argued that this would have little value in itself. Furthermore, some communicators questioned the need to be knowledgeable about species for being able to enjoy, value, or grow interest and insight in nature.

Finally, in 28.7% of the coded answer fragments, communicators compared knowledge about species to things that they attached equal or more importance to, such as interest in and experience of nature, and enjoyment of nature. In particular, communicators stressed the importance of respect and care for nature and species, which they argued should be prioritized. They expressed that as long as people appreciate and cherish nature, knowing much is not really vital. Finally, some communicators emphasized that in-depth knowledge about species and skills such as observing were most important. For instance, they stressed the importance of grasping the ‘big picture’ and becoming aware of interdependencies between species and between species and the environment.

## 4. Discussion

### 4.1. Communicators’ understanding of species literacy levels

In order to build stewardship for biodiversity, communication is needed that strikes a chord with the lay public. For this, communicators need to be aware of perceptions present in their target audiences (Bass et al., 2002; Schutte, 2010; Wratten and Hodge, 1999). We explored biodiversity communicators’ awareness of the species literacy level in primary school children, by asking them to estimate the average score that children aged 9-10 would achieve in an identification test comprising native animal species.

The results demonstrated that most communicators were not aware of the species literacy level in primary school children; their estimations varied widely. In particular, many communicators overestimated the level of species literacy. Surprisingly, experience with children as a target group did not correlate with better estimations. The results are in line with previous studies that have reported professionals in other fields to experience difficulty in estimating prior knowledge levels (Dickens et al., 2013; MacAbasco-O’Connell and Fry-Bowers, 2011; Perrenet, 2010; Schutte, 2010).

The mismatch between estimated and actual knowledge levels indicates a barrier to successful communication. Nature educators might currently not be aware that certain species names of common animals are likely to be perceived by children as jargon. As we expect the mismatch to apply to more than just the identification of species (communicators will probably also overestimate what children know about species’ habitat, diet, and behavior), messages may currently be crafted by communicators that will not be understood as intended.

### 4.2. Species literacy as desired and perceived by communicators

To further put the species literacy level in primary school children into perspective, we compared it with the level as desired by biodiversity communicators and we explored the perceived importance attached to species literacy.

Three quarters of the communicators desired the species literacy level in children to be higher than it actually was. Corroborating these results, communicators generally placed importance on species literacy. Remarkably though, views differed as to why knowledge about species would be important. Some communicators expressed that knowledge about species simply should be part of a person’s knowledge base; e.g., it was stated that people should be familiar with the local environment, which links with the idea that knowledge about flora and fauna can provide people with a ‘sense of place and belonging’ (Horwitz et al., 2001; Standish et al., 2013). Most viewed species literacy not as a goal in itself, but rather as a basic step that helps achieve broader understanding, enriches a person’s life by raising interest and well-being, and/or that instills love and respect for nature. These views are in line with reports that knowledge about species can help shift people’s perceptions and raise affinities towards them (Barnett, 2019; Lindemann-Matthies, 2005; Schlegel and Rupf, 2010; Wilson and Tisdell, 2005) and the notion that species names are part of a language that a person needs to communicate successfully and confidently about nature (Magntorn and Helldén, 2005). The role that communicators ascribed to species knowledge as providing people with insights, e.g. making them aware of changes in the environment, and as contributing to nature experiences, may prove vital at a time when nature degradation continues and people are at an increasing risk of losing connections with nature (Miller, 2005; Pauly, 1995; Pyle, 2011; Soga and Gaston, 2018).

We further note that biodiversity communicators did not attach the same level of importance to different components of species literacy. Most importantly, there was disagreement about the value of naming species. Some communicators stated that naming species has little value in itself, despite the fact that previous authors have argued that a name can be a starting point for more meaningful learning and discussion (Magntorn and Helldén, 2005; Ohl et al., 2014). Similarly, although most communicators wished laypeople to care about nature and to understand ‘the big picture’, some questioned the contribution that species literacy can make in this respect and thus seemed unaware of the role attributed by past authors to factual knowledge in allowing people to build understanding, interest, and appreciation; a pathway that has actually been covered extensively in educational literature (Amer, 2006; Weilbacher, 1993) and has been supported by empirical research (Cosquer et al., 2012; Lindemann-Matthies, 2005; Schlegel and Rupf, 2010; Shwartz et al., 2014). In fact, accessible as they are and easy to relate to, species can be tools in helping people grasp complex, abstract concepts like biodiversity, food webs, and ecosystems (Barker and Slingsby, 1998; Orr, 2005).

### 4.3. Future directions

It is important to mention that we focused our study on estimations of average levels of knowledge, i.e. the identification score that an average child would achieve. However, children differ from one another with respect to what they know, and it is questionable whether communication materials calibrated at an average knowledge level will strike a responsive chord with those who are not average (Wals, 1994). When designing a message aimed at primary school children, it may thus be better to calibrate the level below the actual average level, although the needs of children with greater bodies of knowledge should also not be neglected. Future research could explore how best to address heterogeneous audiences when communicating biodiversity.

Moreover, while we studied communicators’ estimations of the knowledge level in primary school children, future projects could explore the extent to which communicators are aware of perceptions in high school students and adults. For instance, studies could investigate whether communicators working at nature conservancy organizations are aware of knowledge levels in their lay members.

### 4.4. Conclusion

To increase awareness about biodiversity effectively, biodiversity communicators should have a clear picture of prior knowledge in their audiences and the desired outcomes that they strive for. Only then will they be able to meaningfully connect to people’s perceptions and take the necessary steps to achieve the desired level. To our knowledge, this study was the first to investigate species knowledge levels as estimated and desired by biodiversity communicators. We demonstrated that estimating prior knowledge levels in primary school children is difficult for people who communicate about biodiversity, extending the findings in other disciplines (Bass et al., 2002; Kelly and Haidet, 2007; Perrenet, 2010; Storm, 2012). Communicators overestimated and wished for higher knowledge levels in children, suggesting that current educational materials and messages may not connect to existing knowledge. Such misfit between estimated and actual knowledge levels may prevent learning goals from being achieved and may partly explain why conservationists have yet been unsuccessful at reaching certain segments of society.

Moreover, although most biodiversity communicators agreed that species literacy is valuable, we uncovered disagreement among biodiversity communicators as to why species literacy or components of species literacy would be important. This suggests that professionals may benefit from a detailed framework of species literacy that integrates different aspects and values. Such a framework may also encourage biodiversity communicators, educators, and conservationists in their work and could assist them in the design of educational materials and in accounting for the relevance of their activities to society and employers.

Our study further highlights the potential of assessments to bridge the gap between expected and actual knowledge levels (Hailikari et al., 2007). Assessments may help communicators in attuning messages to the appropriate level, in identifying misconceptions to be addressed, and in determining the specific target group that will benefit most from communication or education (Penn et al., 2018; Peterson et al., 2008; Vincenot et al., 2015). Communicators could, for instance, use a series of online quizzes, which would simultaneously provide valuable insights into people’s perceptions, while entertaining participants and encouraging them to learn and find out more about biodiversity, adding to their impact and scope. While we focused on prior knowledge, we recommend that factors such as interest, expectations, and personal experiences are also explored further via such assessments, as they too influence the way people respond to messages, and providing information at the right level will in itself not be enough to change attitudes and behavior (Buijs et al., 2008; Falk and Adelman, 2003; Fischer and Young, 2007; Novacek, 2008; Vázquez-Plass and Wunderle, 2010). As perceptions depend on context and change over time, we recommend assessments to be repeated regularly.

All in all, we demonstrated gaps between the perceived, desired and actual average species literacy level in Dutch primary school children. This suggests that to reach desired knowledge levels in young generations, communicators will benefit from first becoming more aware of current perceptions in children. Efforts to identify, differentiate and get to know the audiences they try to reach would provide biodiversity communicators with opportunities to improve their outreach, which could help achieve broad-based support for conservation.

## Supporting information

Appendix_A_Questionnaire

Appendix_B_Datasheet

## Acknowledgements

We are grateful for the time and effort of all participants in the research. We thank Daniel Oberski for the feedback that we received to improve the survey.

## Role of the funding source

The study did not receive any specific grant from funding agencies in the public, commercial, or not-for-profit-sectors.

## Declaration of competing interest

We have no conflicts of interest to disclose.

## Notes

### Competing Interest Statement

The authors have declared no competing interest.

