## Appendix A for "Children’s Species Literacy as Estimated and Desired by Biodiversity Communicators: a Mismatch with the Actual Level"

### APPENDIX A – Questionnaire

This Word-file contains the questionnaire used for targeting biodiversity communicators. The survey was made in Qualtrics (https://www.qualtrics.com) and distributed online.

### Questionnaire (English translation)

#### *Species Knowledge in the Netherlands – NAME INSTITUTE*

How well can you recognize animal species? And how well do you think others can do that? With this quiz you can test your species knowledge yourself ánd you help us with our research on knowledge about nature in the Netherlands. The quiz will take approximately 10 minutes and your answers will be processed anonymously by [NAME DEPARTMENT]. There are no wrong answers! Answer everything as honestly as possible and don't look for answers via Google.

Thanks in advance!

Do you have any questions or remarks? You can send these to [NAMES].

If you continue, you consent to anonymous processing of your answers by [NAME DEPARTMENT].

1) Do you do anything with biodiversity, nature and/or animals in your (voluntary) work?

- Yes

- No

- I am an (intern) student

2) Do you also do something with communication in your work? That is: do you convey information to a public about biodiversity, nature and/or animals?

- Yes

- No

**[Block Background biodiversity and communication]**

(*These questions are only displayed to participants who have (voluntary) work related to biodiversity and who communicate about it to an audience)*

3) We would like to find out some things about your background via the questions below.

You have indicated that in your position you do something with biodiversity and communication. Can you briefly elaborate on this? (e.g. verbally through presentations, written texts in a magazine, guided tours, website, blog, TV, films, radio...)

4) Which organization(s) do you work for?

5) What is your highest achieved level of education?
- WO (research university)
- HBO (university of applied sciences)
- MBO (vocational training)
- Secondary school
- Different

6) Have you followed education related to nature or biodiversity?
- Yes
- No

7) In the past five years, has the general public been (one of) your target audiences?
- Yes
- No

8) In the past five years, have primary school children aged 9-10 been (one of) your target audiences?
- Yes
- No

**[Block No biodiversity]**

*(These questions were shown to participants that indicated that they did not do (voluntary) work on biodiversity and communication)*

9) What is your highest achieved level of education?
- WO (research university)
- HBO (university of applied sciences)
- MBO (vocational training)
- Secondary school
- Different

10) In which field do you work?

11) Which study/studies do you do?

12) Are you a member of (a) nature organization(s)? If yes, which one?

- No

- Yes, I am member of... (fill in below)

____________________________________

**[Block Identifcation test]**

The species-quiz starts here. You will see 27 animal species. Type in the name. It is okay if you do not know a species or if you fill in an incorrect answer, because the purpose of this research is to see how much people currently know. It is therefore important not to use tools such as Google to find answers and not to ask colleagues.

When providing answers, be as specific as possible. For example, you may not know that the photo below depicts a lion, but you may know that it is a mammal, a carnivoran or a feline. If you really don't know, you can enter "I don't know" or a question mark.

The answers are displayed at the very end of the test.

*Have fun!*

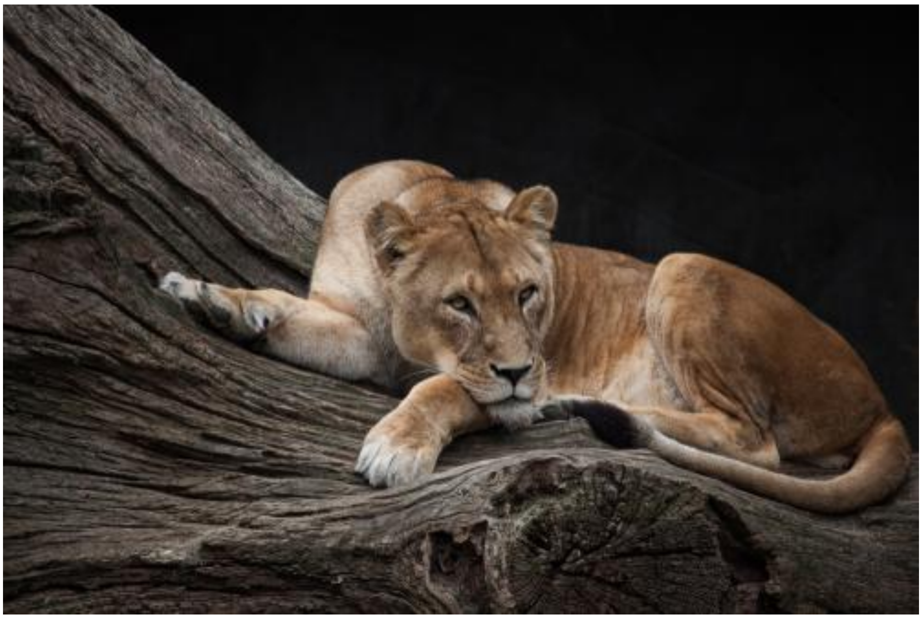

*The quiz starts below.*

[Questions 13 to 39 are the questions that belong to the species identification test. See page 6 for the animals that were successively presented; for each animal the question was asked: "What is the name of this animal?").]

---

40) How many of these animals do you think you have identified correctly? (slider-question of 0-27)

**[Block: Estimations]**

41) How do you estimate the species knowledge of others? Suppose that the two groups below were to take part in the species-quiz. How many animals would they name correctly?

General public (slider-question of 0-27)
Primary school children at fourth grade level (aged 9-10) (slider-question of 0-27)

42) How many animals do you think the following two groups should be able to name correctly?

General public (slider-question of 0-27)
Primary school children at fourth grade level (aged 9-10) (slider-question of 0-27)

**[Block: Importance]**

In this section you will see a number of statements. Please indicate to what extent you agree with the statements. It would help us if you could elaborate on your choice. If you do not know what you think, or if you do not have an opinion, please indicate this in the explanation.

43) It is important that people are interested in nature. (scale of 0 – 10)

Remarks:

__________________________________________________________________

__________________________________________________________________

44) It is important that people recognize many different animal species. (scale of 0 – 10)

Remarks:

__________________________________________________________________

__________________________________________________________________

45) In primary education, attention must be paid to species knowledge. (scale of 0 – 10)

Remarks:

__________________________________________________________________

__________________________________________________________________

46) Species knowledge stimulates interest in nature. (scale of 0 – 10)

Remarks:

__________________________________________________________________

__________________________________________________________________

**[Block: Tot slot]**

Is there anything else you would like to note?

__________________________________________________________________

__________________________________________________________________

Do you have any feedback about this questionnaire?

__________________________________________________________________

__________________________________________________________________

Are you curious about the results of our research? If you leave your e-mail address below, we will send you the final results (once).

__________________________________________________________________

### Animal Species in the Identification Test

| | Species |
|---|---|
| 1 | *Vulpes vulpes* |
| 2 | *Chloris chloris* |
| 3 | *Meles meles* |
| 4 | *Erithacus rubecula* |
| 5 | *Vanessa atalanta* |
| 6 | *Coloeus monedula* |
| 7 | *Erinaceus europaeus* |
| 8 | *Turdus merula* |
| 9 | *Bufo bufo* |
| 10 | *Araneus diadematus* |
| 11 | *Gallinula chloropus* |
| 12 | *Lepus europaeus* |
| 13 | *Pica pica* |
| 14 | *Aglais urticae* |
| 15 | *Fringilla coelebs* |
| 16 | *Sus scrofa* |
| 17 | *Porcellio scaber* |
| 18 | *Cyanistes caeruleus* |
| 19 | *Aegithalos caudatus* |
| 20 | *Sciurus vulgaris* |
| 21 | *Alcedo atthis* |
| 22 | *Capreolus capreolus* |
| 23 | *Limosa limosa* |
| 24 | *Canis lupus* |
| 25 | *Podiceps cristatus* |
| 26 | *Passer domesticus* |
| 27 | *Lutra lutra* |

### Pictures used in the Identification Test

All pictures were freely downloaded from the website https://pixabay.com/ (Pixabay License)

| | |
|---|---|
| 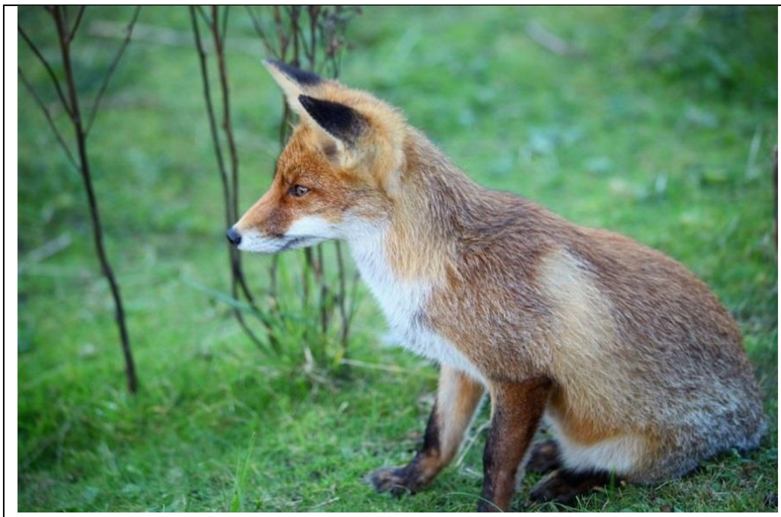 | Species:<br>*Vulpes vulpes*<br><br>Source:<br>https://pixabay.com/nl/fox-dierlijke-wild-2066027/<br><br>Pixabay-user: gabrielgs |
| 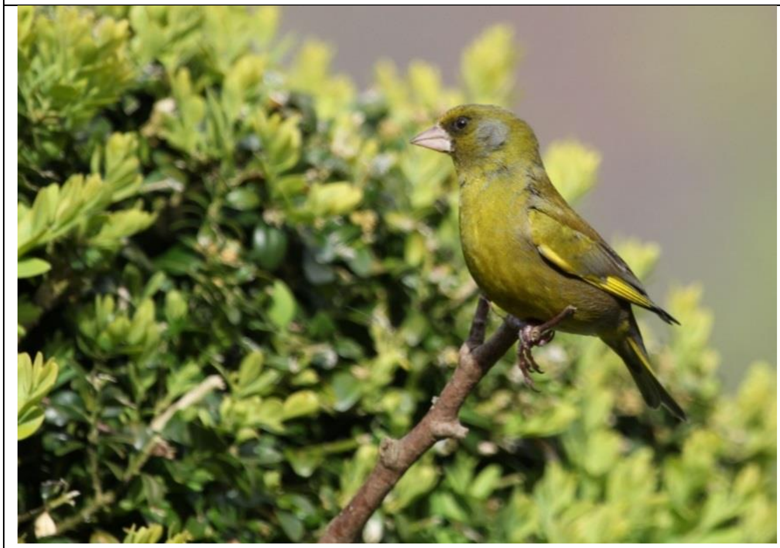 | Species:<br>*Chloris chloris*<br><br>Source:<br>https://pixabay.com/nl/vogel-finch-groene-zitten-veer-113496/<br><br>Pixabay-user: ginger |
| 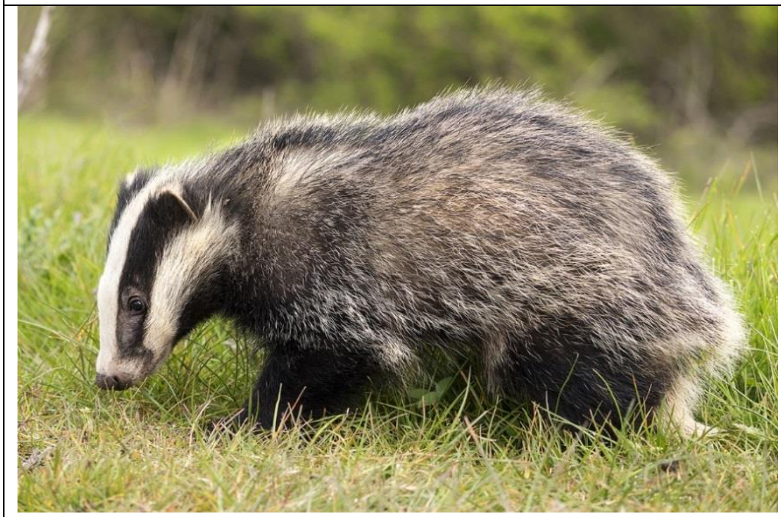 | Species:<br>*Meles meles*<br><br>Source:<br>https://pixabay.com/nl/badger-brock-dierlijke-zoogdier-2030980/<br><br>Pixabay-user: andyballard (Andy Ballard) |

| | |
|---|---|
| 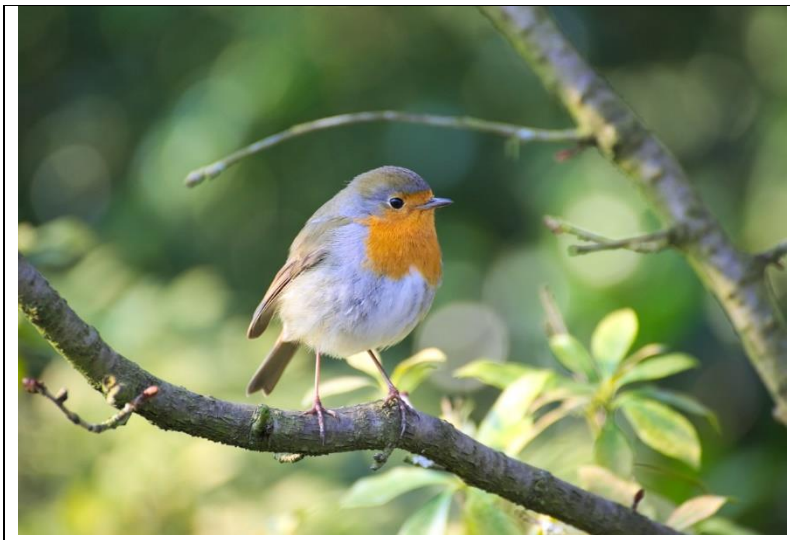 | Species:<br>*Erithacus rubecula*<br><br>Source:<br>https://pixabay.com/nl/robin-vogel-tuin-erithacus-rubecula-2187366/<br><br>Pixabay-user: EvgeniT (Evgeni Tcherkasski) |
| 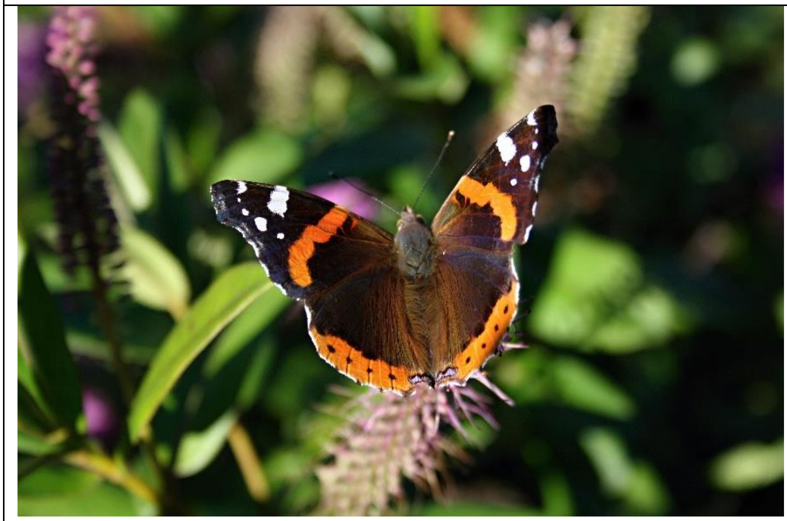 | Species:<br>*Vanessa atalanta*<br><br>Source:<br>https://pixabay.com/nl/vlinder-vanessa-atalanta-red-admiral-2408696/<br><br>Pixabay-user: ctaklis (Χρήστος Τακλής) |
| 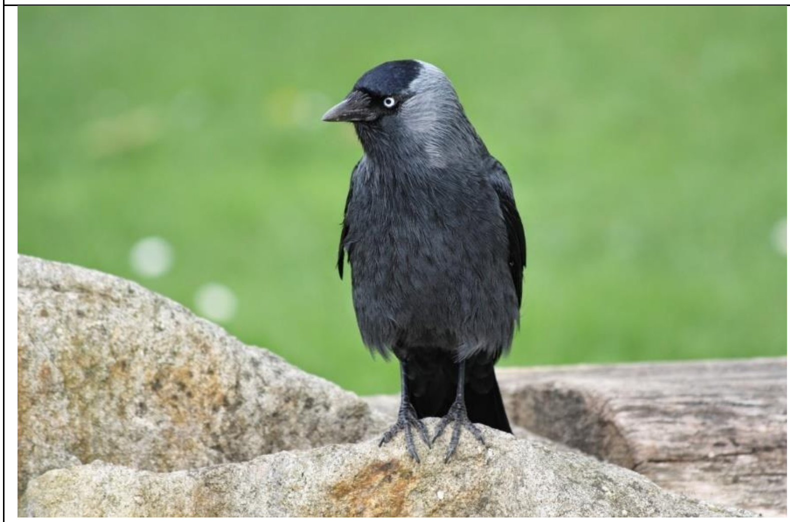 | Species:<br>*Coloeus monedula*<br><br>Source:<br>https://pixabay.com/nl/vogel-kauw-zwart-blue-eye-1268303/<br><br>Pixabay-user: DocChicago |

| | |
|---|---|
| 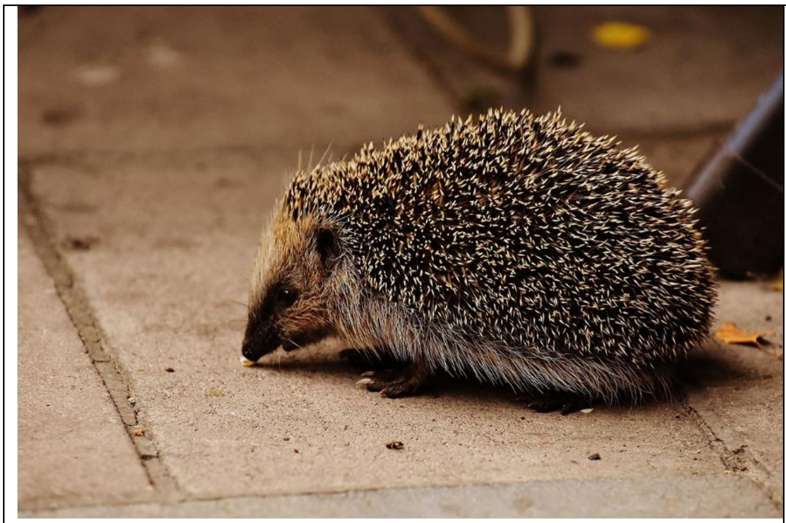 | Species:<br>*Erinaceus europaeus*<br><br>Source:<br>https://pixabay.com/nl/egel-kind-jonge-egel-egel-dierlijke-1758877/<br><br>Pixabay-user: Alexas_Fotos (Alexandra) |
| 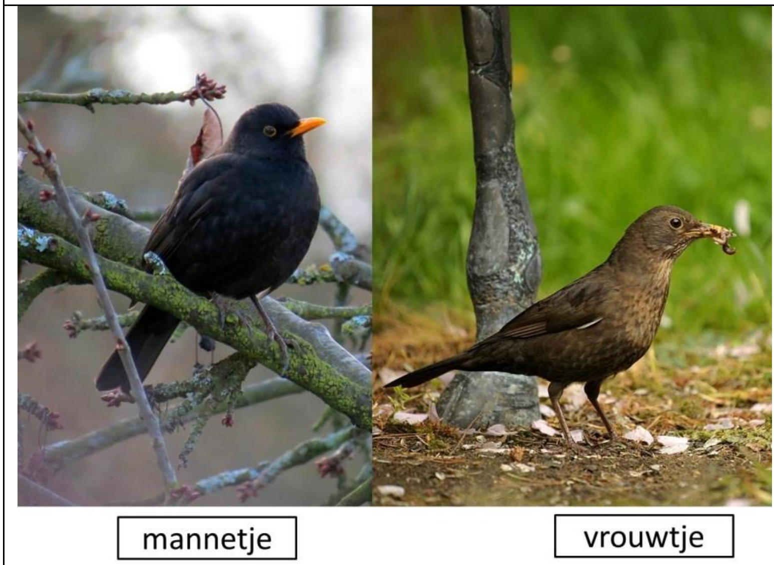  | Species:<br>*Turdus merula*<br>Source (male):<br>https://pixabay.com/nl/blackbird-vogel-de-winter-kerseboom-1960706/<br>Pixabay-user: manfredrichter (Manfred Richter)<br><br>Source (female):<br>https://pixabay.com/nl/blackbird-vogel-songbird-tuin-1758790/<br>Pixabay-user: susannp4 (Sussan Mielke) |
| 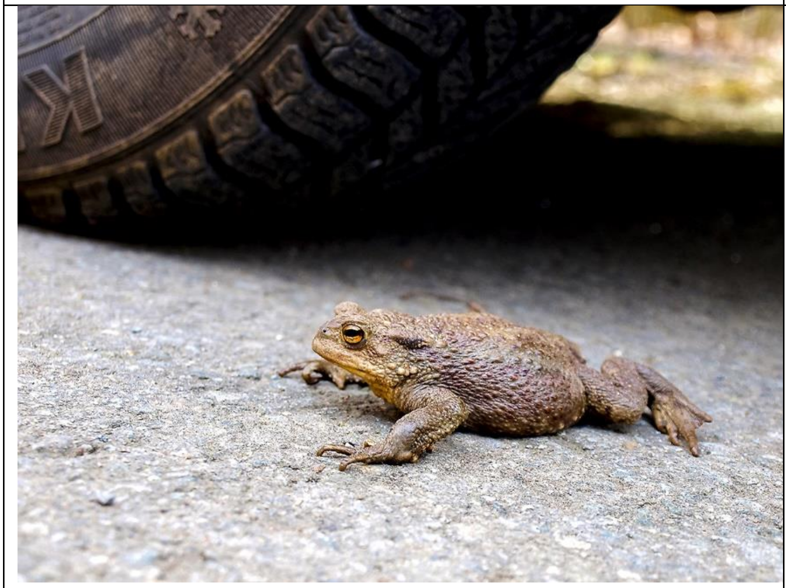 | Species:<br>*Bufo bufo*<br><br>Source:<br>https://pixabay.com/nl/gewone-pad-toad-amfibie%C3%ABn-de-natuur-2382964/<br>Pixabay-user: Kathy2408 (Kathy Büscher) |

| | |
|---|---|
| 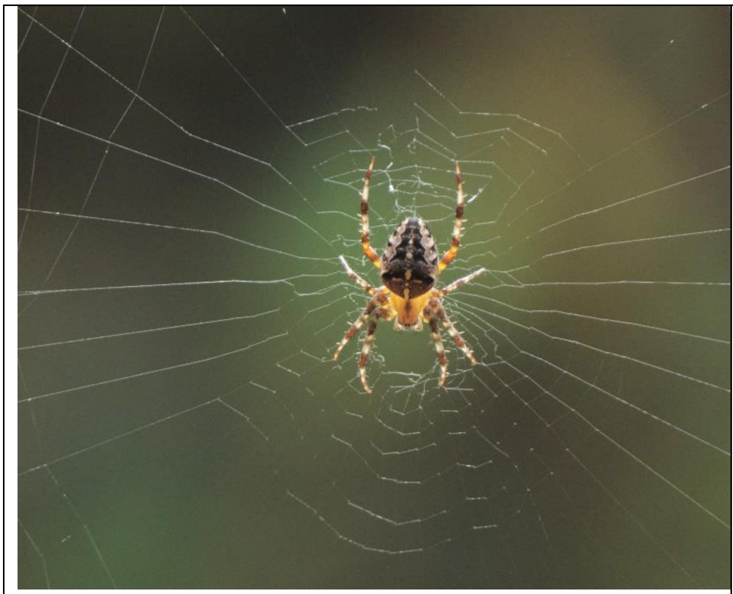 | Species:<br>*Araneus diadematus*<br><br>Source:<br>[https://pixabay.com/nl/users/timspitzer-5662946/](https://pixabay.com/nl/users/timspitzer-5662946/)<br><br>Pixabay-user: timspitzer (Tim Spitzer) |
| 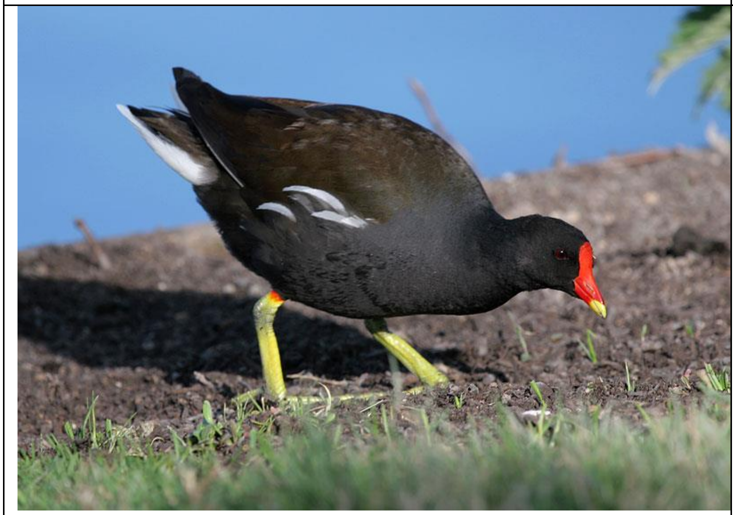 | Species:<br>*Gallinula chloropus*<br><br>Source:<br>https://commons.wikimedia.org/wiki/File:Kokoszka(Grzecho_Lukasik).jpg<br><br>Assumed Author: G. Lukasik |
| 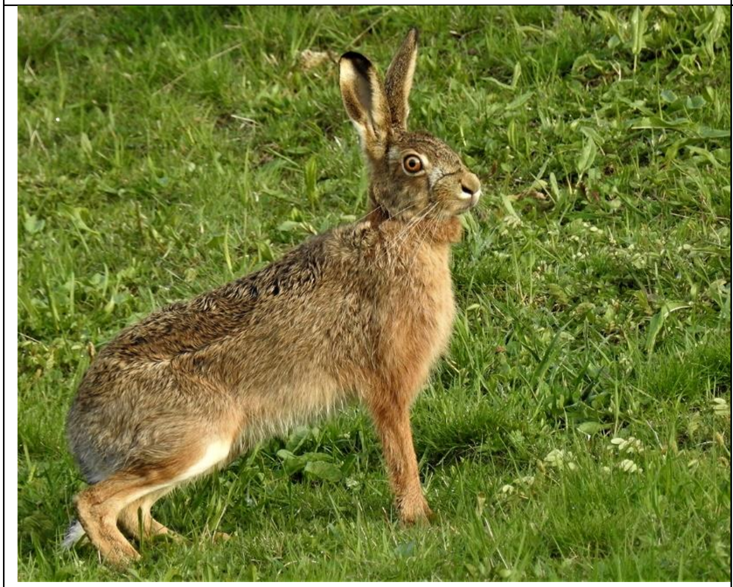 | Species:<br>*Lepus europaeus*<br><br>Source:<br>[https://pixabay.com/nl/haas-lepus-europaeus-2240059/](https://pixabay.com/nl/haas-lepus-europaeus-2240059/)<br><br>Pixabay-user:<br>vetler |

| | |
|---|---|
| 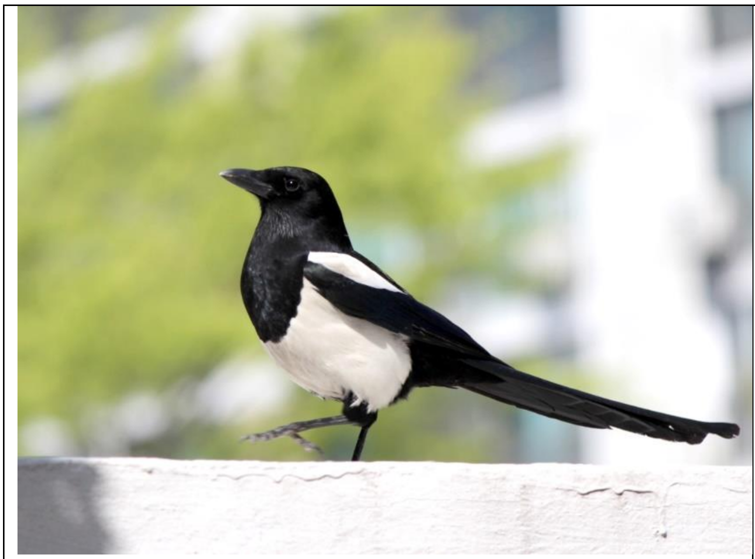 | Species:<br>*Pica pica*<br><br>Source:<br>https://pixabay.com/nl/ekster-nieuw-vogels-481960/<br><br>Pixabay-user: jinhokim (Jinho Kim) |
| 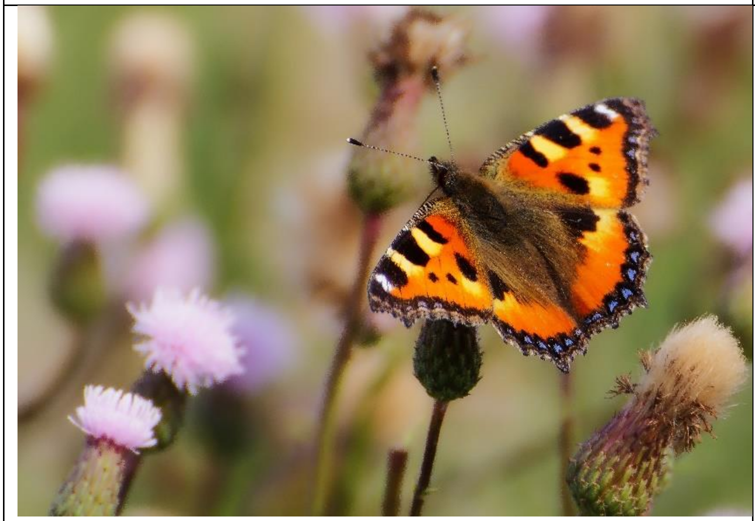 | Species:<br>*Aglais urticae*<br><br>Source:<br>https://pixabay.com/nl/dieren-vlinder-kleine-vos-insect-176860/<br><br>Pixabay-user: Kalahari |
| 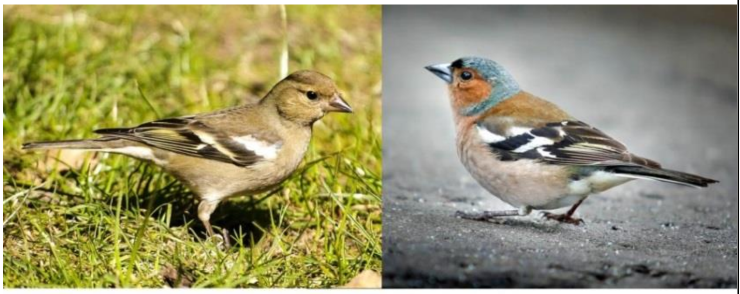<br>vrouwtje mannetje | Species:<br>*Fringilla coelebs*<br>Source (female):<br>https://pixabay.com/nl/denunciateur-vink-vogel-vogel-2404776/<br>Pixabay-user: Kathy2408 (Kathy Büscher)<br><br>Source (male):<br>https://pixabay.com/nl/vink-vogel-mus-park-%C3%A9%C3%A9n-2491484/<br>Pixabay-user: klimkin (svklimkin) |

| | |
|---|---|
| 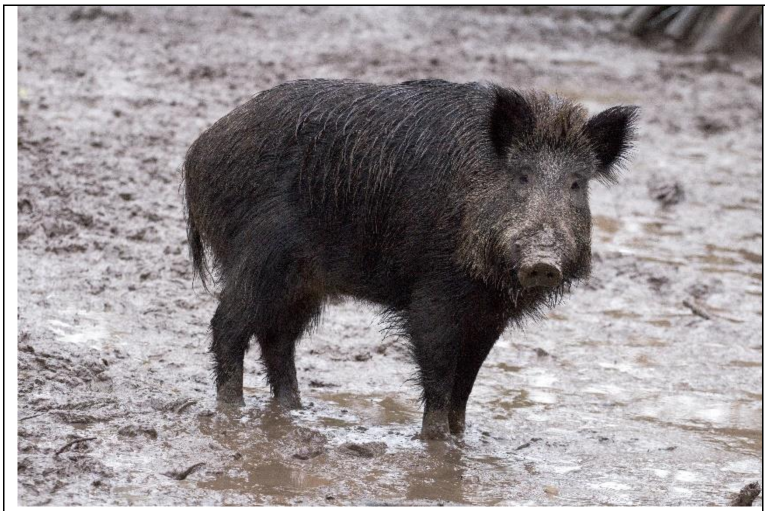 | Species:<br>*Sus scrofa*<br><br>Source:<br>[https://pixabay.com/nl/wilde-wilde-zwijnen-bos-wild-zwijn-659379/](https://pixabay.com/nl/wilde-wilde-zwijnen-bos-wild-zwijn-659379/)<br>Pixabay-user: domeckopol (Andreas N) |
| 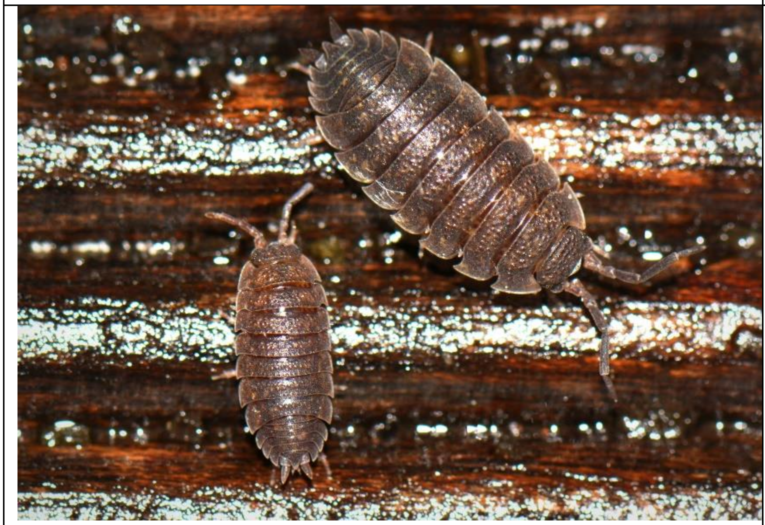 | Species:<br>*Porcellio scaber*<br><br>Source:<br>[https://pixabay.com/nl/insecten-dieren-macro-pissebed-292482/](https://pixabay.com/nl/insecten-dieren-macro-pissebed-292482/)<br><br>Pixabay-user: Ben_Kerckx (Ben Kerckx) |
| 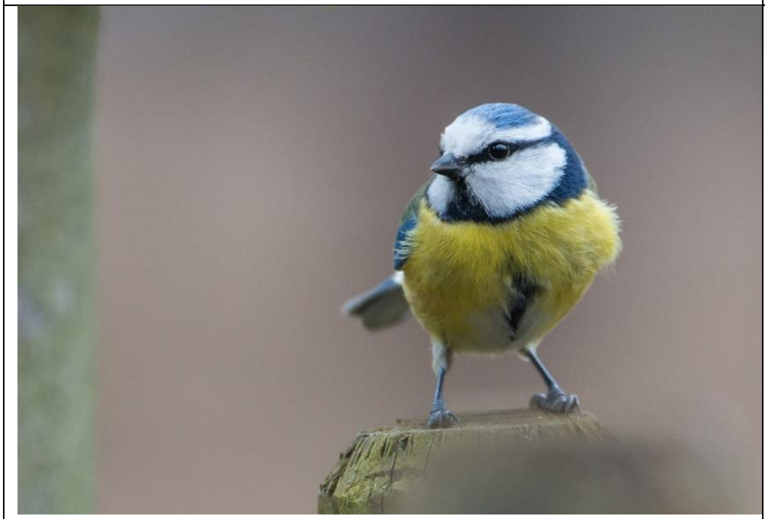 | Species:<br>*Cyanistes caeruleus*<br><br>Source:<br>[https://pixabay.com/nl/pimpelmees-vogel-paridae-2109987/](https://pixabay.com/nl/pimpelmees-vogel-paridae-2109987/)<br><br>Pixabay-user: wolfgang_vogt (Wolfgang Vogt) |

| | |
|---|---|
| 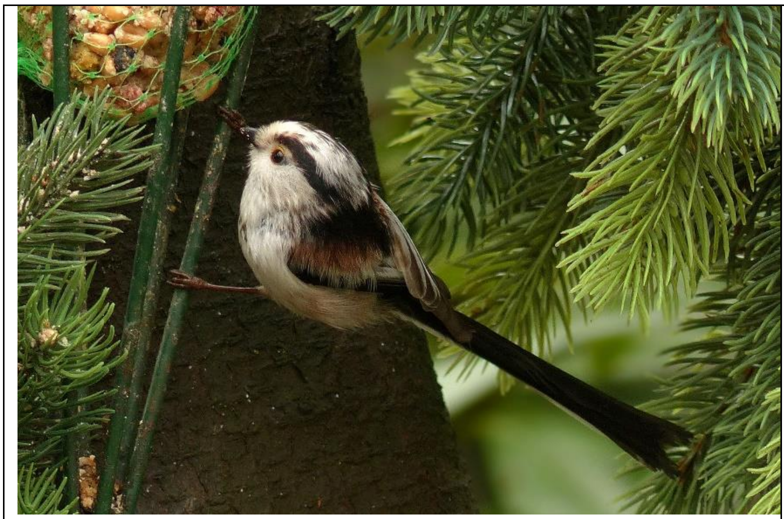 | Species:<br>*Aegithalos caudatus*<br><br>Source:<br>https://pixabay.com/nl/vogel-staartmees-foerageren-tuin-675131/<br>Pixabay-user: Oldiefan (Christiane) |
| 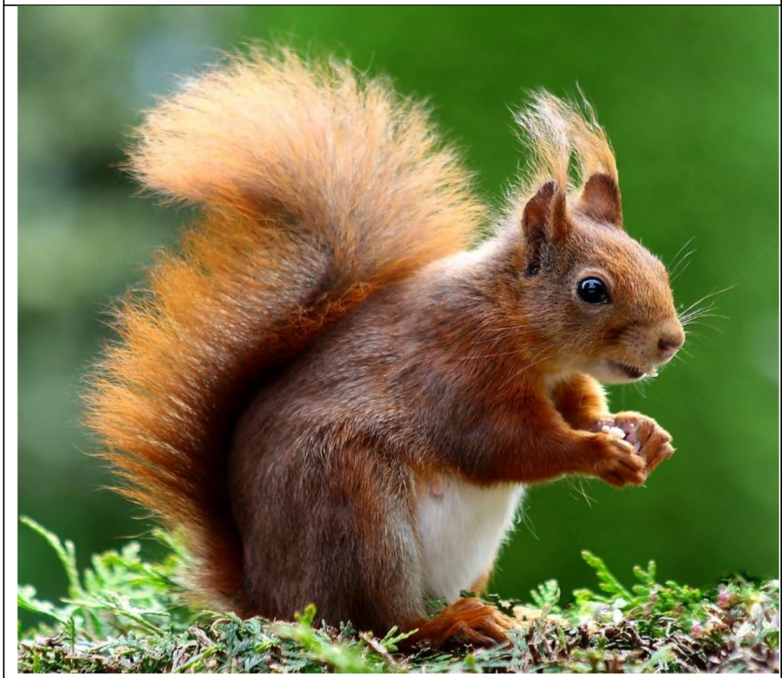 | Species:<br>*Sciurus vulgaris*<br><br>Source:<br>https://pixabay.com/nl/eekhoorn-dierlijke-cute-knaagdieren-493790/<br><br>Pixabay-user: Elli60 (Elli Stattaus) |
| 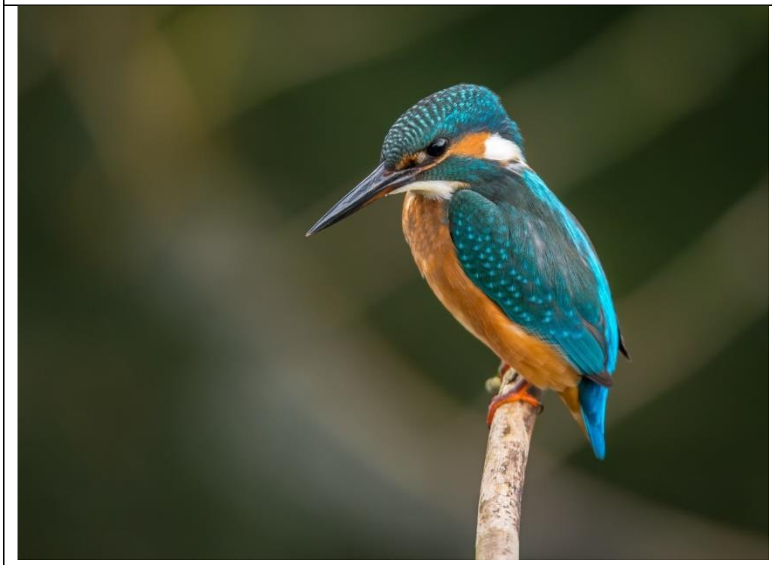 | Species:<br>*Alcedo atthis*<br><br>Source:<br>https://pixabay.com/nl/ijsvogel-vogel-dierlijke-983944/#<br>Pixabay-user: Free-Photos |

| | |
|---|---|
| 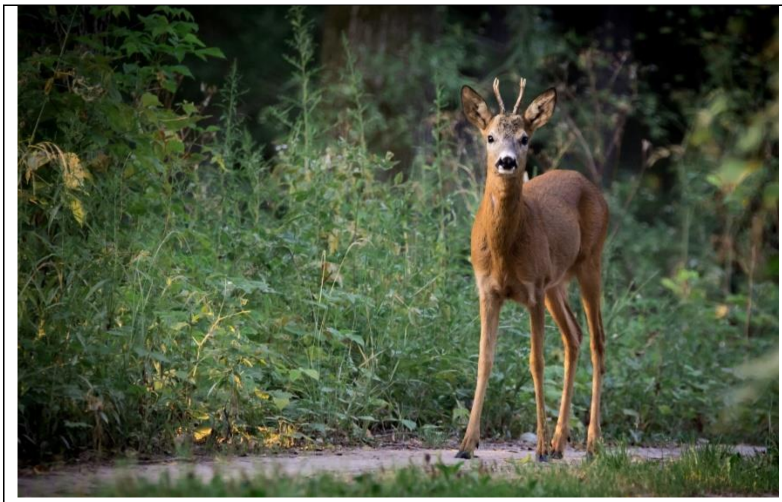 | Species:<br>*Capreolus capreolus*<br><br>Source:<br>https://pixabay.com/nl/ree-capreolus-capreolus-hinde-dier-880581/<br>Pixabay-user: LubosHouska (Lubos Houska) |
| 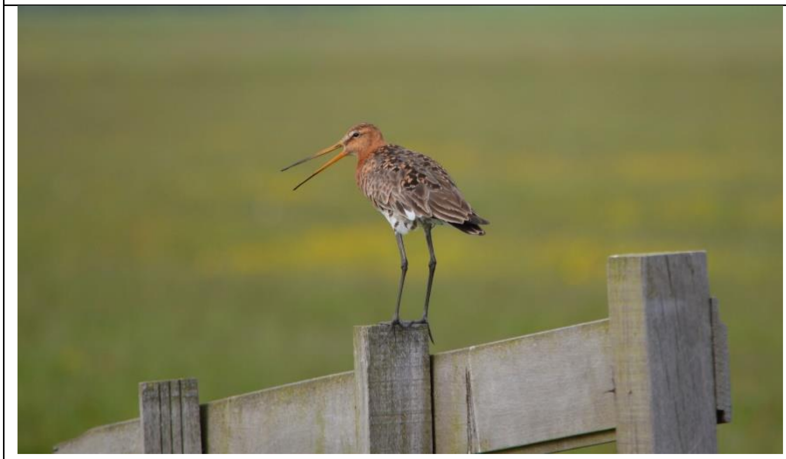 | Species:<br>*Limosa limosa*<br><br>Source:<br>https://pixabay.com/nl/godwit-paal-vogel-natuur-dierlijke-164593/<br><br>Pixabay-user: PublicDomainPictures |
| 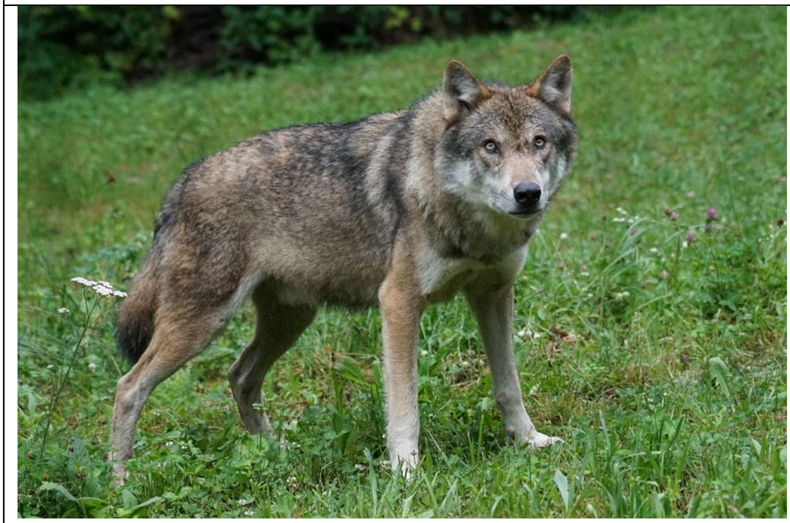 | Species:<br>*Canis lupus*<br><br>Source:<br>https://pixabay.com/nl/wolf-roofdier-europese-wolf-1514782/<br>Pixabay-user: Pixel-mixer (Marcel Langthim) |

| | |
|---|---|
| 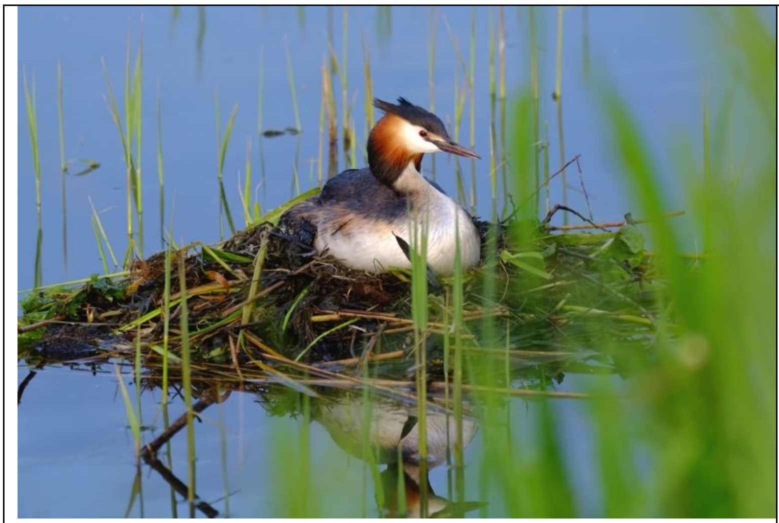 | Species:<br>*Podiceps cristatus*<br><br>Source:<br>https://pixabay.com/nl/fuut-vogel-watervogel-meer-water-2461262/<br>Pixabay-user: doreen_kinistino (Doreen Sawitza) |
| 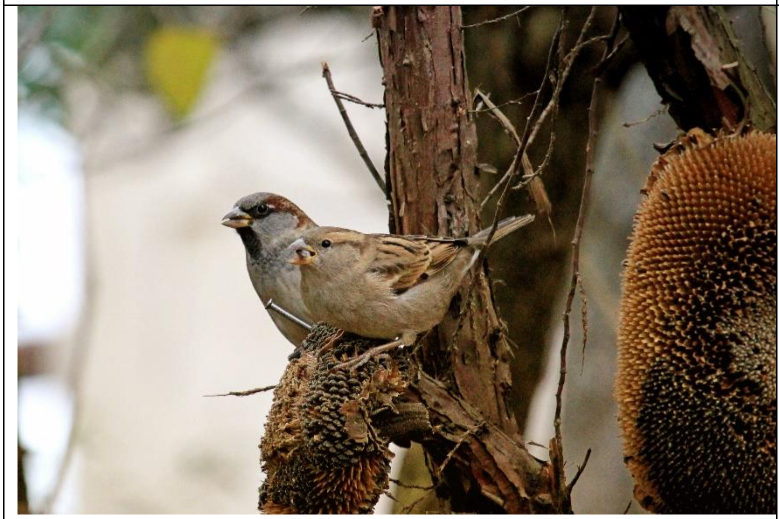 | Species:<br>*Passer domesticus*<br><br>Source:<br>https://pixabay.com/nl/veevoeder-boom-vogel-mus-herfst-1749194/<br>Pixabay-user: hansbenn (Hans Benn) |
| 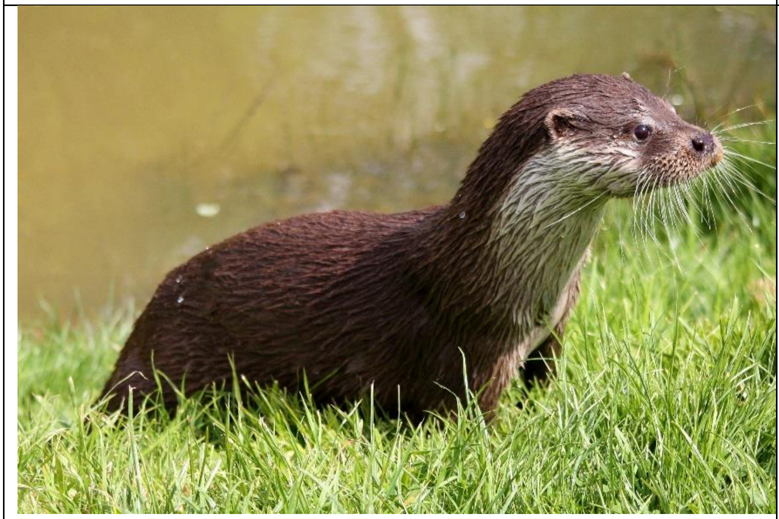 | Species:<br>*Lutra lutra*<br><br>Source:<br>https://pixabay.com/nl/otter-het-wild-levende-dieren-natuur-1914239/<br>Pixabay-user: little_arrows (Peter Hoare) |
